# Label-Free Determination of Chondroitin Sulphate from Microgram Quantities of Human Milk

**DOI:** 10.64898/2026.05.08.723732

**Authors:** Melissa Greenwood, Sean Austin, Patricia Murciano-Martinez, Katherine A. Hollywood, Marina Machidon, Reynard Spiess, Janet Berrington, Sabine Flitsch, Perdita Barran, Christopher J Stewart

## Abstract

Human milk contains structurally diverse glycans with key roles in shaping infant development, yet analytical constraints limit characterisation from low‐volume samples. Glycosaminoglycans (GAGs), including chondroitin sulphate (CS), are understudied due to existing protocols requiring sample volumes of at least 5 mL and lengthy extraction steps prior to instrumental analysis. This study establishes a workflow for quantifying CS disaccharides from 25 µL of human milk, enabling analysis of samples previously inaccessible to GAG profiling, such as those collected as salvage samples from neonatal intensive care units. For CS quantification, the CS is first enzymatically depolymerised using chondroitinase ABC to release repeating disaccharide units. Matrix complexity is reduced *via* two rounds of acetonitrile‐based protein and lipid precipitation. Disaccharides are separated by hydrophilic interaction liquid chromatography and detected using a Triple Quadrupole Mass Spectrometer, providing robust sensitivity for all CS disaccharides. Method development and validation were performed using pooled mature human milk from term infants. This workflow facilitates detection of all CS disaccharides, with low but reproducible recoveries for total CS. Low‐ and high‐level spike recoveries were 41.3% (RSD_r_ 7.5%, RSD_iR_ 15.9%) and 43.7% (RSD_r_ 24.4%, RSD_iR_ 27.9%), respectively. Despite modest absolute accuracy, precision remained sufficient to make relative comparison of CS concentrations between samples. This method expands the analytical toolkit for human milk glycomics, enabling same day preparation and CS profiling from sample volumes that are 200 times smaller than prior work, supporting future investigations into GAG‐mediated functions in early life.

Graphical abstract
Schematic of sample preparation protocol
25 μL of human milk is combined with lyase enzymes and TRIS buffer containing the internal standard prior to incubation. Samples then undergo multiple rounds of centrifugation and refrigeration before analysis *via* LC-MS/MS. Made using BioRender.com. Glycan nomenclature following Varki et al., 2015.

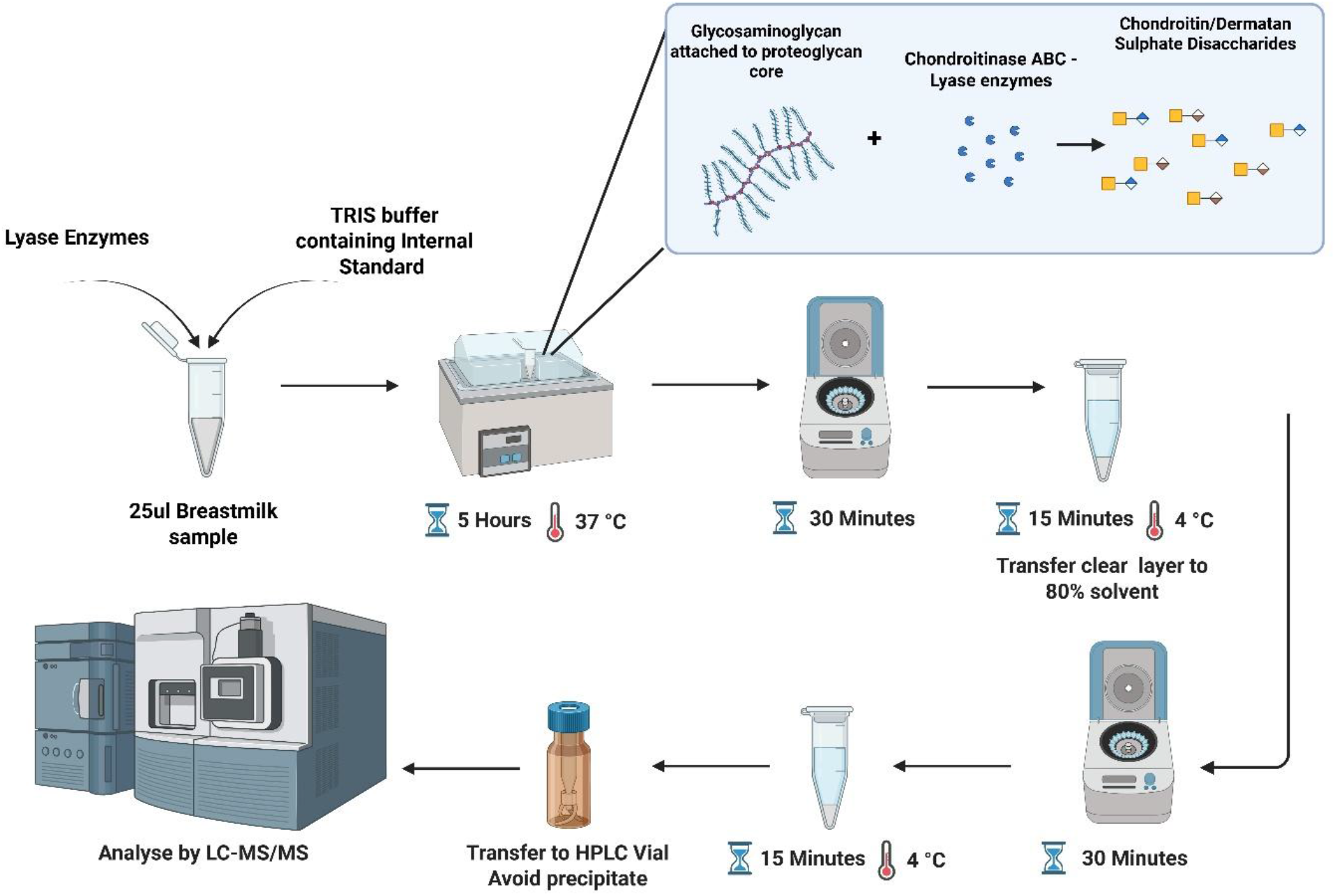

## INTRODUCTION

Human milk is a complex biofluid enriched with diverse bioactive components that support infant nutrition, immune development, and infant gut microbiota (Masi et al. 2024; Stewart 2023). Its composition is dynamic and influenced by maternal–infant interactions, yet many components remain poorly characterised due to analytical constraints, particularly when only small sample volumes are available (Granger et al. 2021). This limitation is especially relevant for studies involving preterm infants, longitudinal sampling, or biobanked samples, where milk volumes are restricted. Moreover, the ability to perform reliable analyses from low volume samples has become increasingly important in the multi-omics era, in which multiple assays are performed on a single sample and conserving material maximises the range of analyses that can be performed.

Glycosaminoglycans (GAGs) are complex, negatively charged polysaccharides, that are present across the body and recognised as bioactive glycans in human milk. The gut microbiome exhibits glycan utilisation ‘preference’ shaped by the dominant microbial taxa, hence glycans available for consumption can influence which species thrive. Endogenous glycan utilisation has been shown to affect colonic health, highlighting the potential functional relevance of milk-derived GAGs (Koropatkin et al., 2012; Masi et al. 2024). Among the major GAG classes, chondroitin sulphate (CS) is of particular interest due to its structural diversity, variable sulphation and its position as the most abundant GAG in human milk (Figure 1) (Casale 2023; Coppa et al. 2013; Szekeres et al., 2022). These features determine biological function but also create analytical challenges, as some CS disaccharides are isomeric and require high‐resolution separation and sensitive detection (Szekeres et al., 2022).

**Figure 1:**
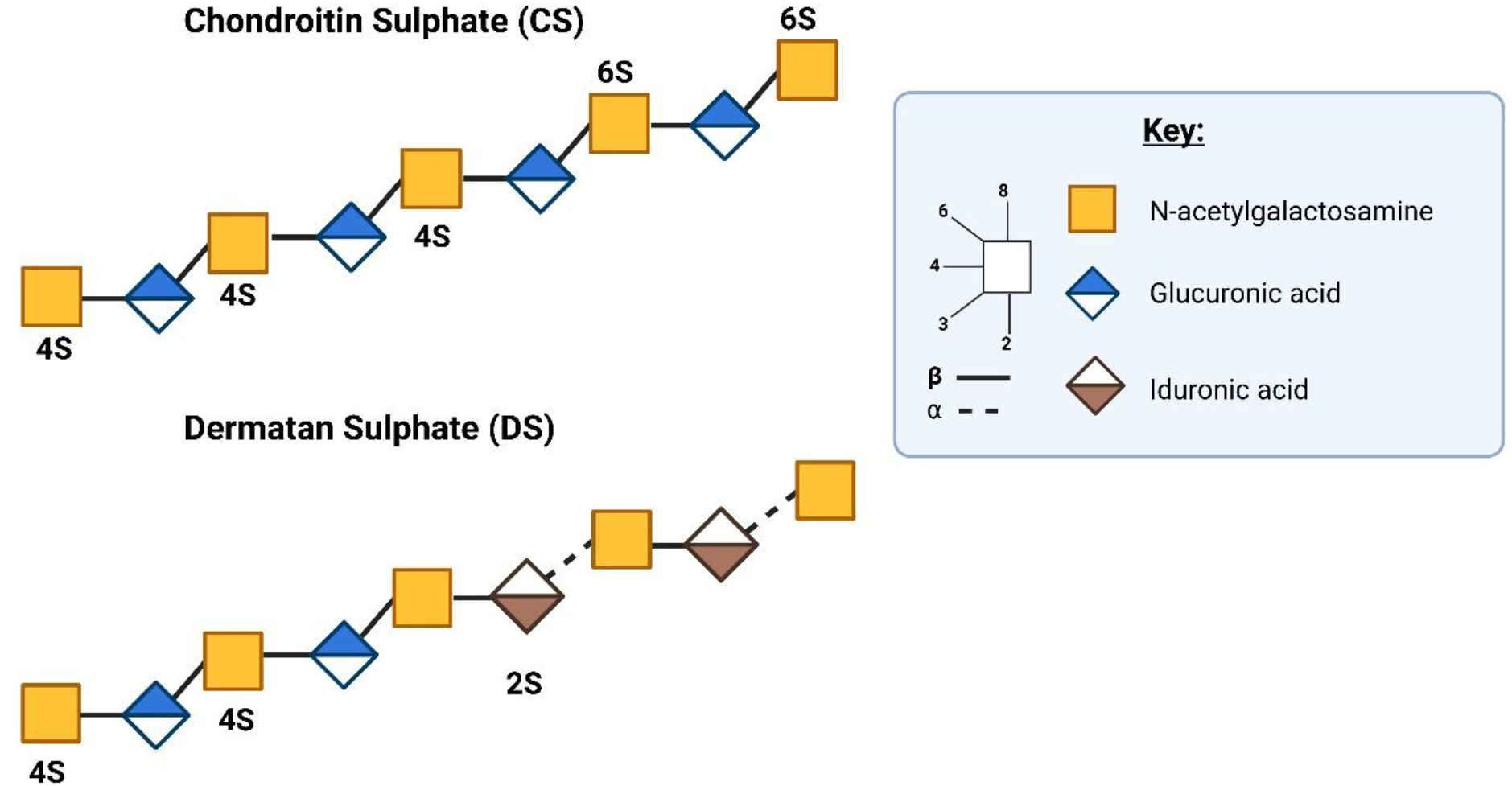
Polysaccharide structures of CS and dermatan sulphate (DS) chains (Greenwood et al., 2024). Made using Biorender.com. Glycan nomenclature following Varki et al., 2015. CS and other GAGs are typically analysed after enzymatic depolymerisation, in which bacterial lyases cleave polysaccharide chains into defined disaccharide units (Casale 2023; Linhardt 2001). Detection can be further enhanced by application of fluorescent tags; most commonly 2-aminoacridone (2-AMAC), which attaches to the reducing end of the disaccharide which increases hydrophobicity and enables detection by UV or fluorescence (FLD) (Vreeker and Wuhrer 2017). This approach provides structural resolution but demands analytical workflows capable of handling highly charged, sulphated species with overlapping retention and fragmentation profiles. As a result, robust quantification of CS disaccharides in complex matrices such as human milk remains technically demanding, particularly when sample volumes are limited.

Multiple analytical approaches have been used for GAG quantification, including carbazole assays, agarose-gel electrophoresis, and liquid chromatography (LC) (Masi et al., 2024). LC coupled with tandem mass spectrometry (LC-MS/MS) offers the greatest sensitivity and structural resolution for sulphated disaccharides (Volpi 2010; Volpi et al., 2014; Coppa et al., 2013). LC–MS/MS enables separation of isomeric GAG disaccharides and sensitive precursor– product ion detection, making it the current standard approach for detailed GAG profiling.

Existing workflows for biological matrices, such as those developed by Volpi et al., rely on multi-step sample preparation involving proteolysis, centrifugal filtration, enzymatic depolymerisation, and fluorescent labelling prior to reverse phase liquid chromatography (RP-LC) on-line with FLD and electrospray ionization (ESI) MS (Volpi et al., 2014; Volpi 2010). Adaptations of these methods have been applied to human milk, most notably by Coppa et al., and Wang et al., who used multi‐day protocols requiring 5–10 mL of sample (Coppa et al. 2011; Coppa et al. 2010; Wang et al. 2018; Volpi 2010). These studies demonstrated the presence of CS, DS, and heparin/heparan sulphate in human milk but highlighted a major limitation: current methods depend on large sample volumes and lengthy preparation, restricting their use in studies where only small volumes are available.

## RESULTS AND DISCUSSION

### Early Method Development for the Detection of CS Disaccharides in Human Milk

RP-LC based detection of AMAC labelled GAGs, as described by Volpi et al., (2010) was insufficiently sensitive for detection of endogenous human milk GAGs, with AMAC labelled CS disaccharide standards presenting significant problems with chromatographic peak resolution and peak splitting. Likewise, AMAC labelled disaccharides detected using hydrophilic interaction chromatography (HILIC) was unsuccessful. Therefore, 2-Aminobenzamide (2-AB) labelling and HILIC chromatography was implemented as it is well established for analysing other oligosaccharides (Guile et al., 1996; Austin and Benet 2018) (Figure 2). This method gave excellent chromatographic performance but was insufficiently sensitive for application to GAG analysis in human milk, with enzyme treated human milk showing no detectable disaccharides (Figure 2).

**Figure 2:**
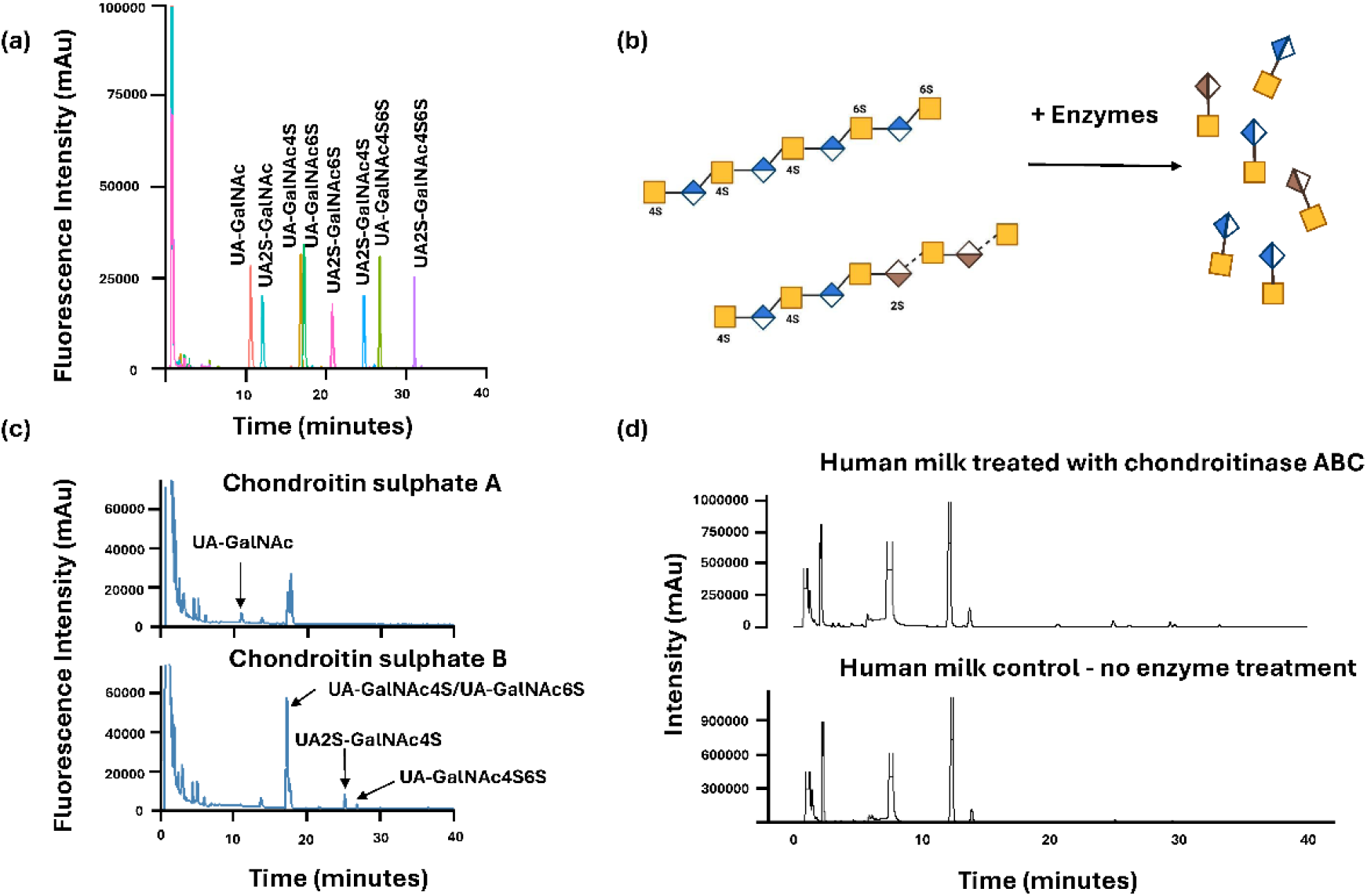
Steps taken in early method development. (a) UHPLC-FLD of 2-AB labelled CS standard disaccharides prepared in water, analysed by HILIC. (b) Schematic demonstrating a simplified enzymatic degradation of CS and DS polysaccharide chains into disaccharide units. (c) Chondroitin sulphate A and chondroitin sulphate B prepared in water, digested with chondroitinase ABC. Disaccharide products labelled with 2-AB and analysed *via* HILIC UPHLC FLD. (d) Human milk treated with chondroitinase ABC ran in parallel with human milk undergoing no enzymatic digestion, demonstrating the insufficiency of FLD-LC alone to detect CS analytes. Samples labelled with 2-AB and analysed *via* HILIC UHPLC FLD. Glycan nomenclature following Varki et al., 2015.

The method was therefore transferred to LC-MS/MS which supported simplified sample preparation, as there was no longer a need to use the 2-AB label for detection. While this simplified the sample preparation, the isomers UA-GalNAc4S and UA-GalNAc6S could no longer be chromatographically separated, however, they could be differentiated using MS/MS since they fragment differently.

### Method Development

Detection of endogenous CS disaccharides from human milk samples required increased sensitivity *via* LC-MS/MS, negating the need for labelling. Partial separation of the seven CS disaccharides and the internal standard was achieved using HILIC on an ACQUITY™ Premier Glycan BEH Amide (130 Å, 1.7 µm, 2.1 × 150 mm) with a gradient of 50mM ammonium formate (pH 4.4) and acetonitrile (Figure 3). Baseline resolution was obtained for UA-GalNAc, UA2S-GalNAc, UA2S-GalNAc6S, UA2S-GalNAc4S, and UA-GalNAc4S6S. However, the isomers UA-GalNAc4S and UA-GalNAc6S co-eluted under all chromatographic conditions tested and therefore required differentiation by their optimised multiple reaction monitoring transitions (MRM). The close elution of UA2S-GalNAc6S and the internal standard also benefited from transition-specific selectivity (Figure 3).

**Figure 3:**
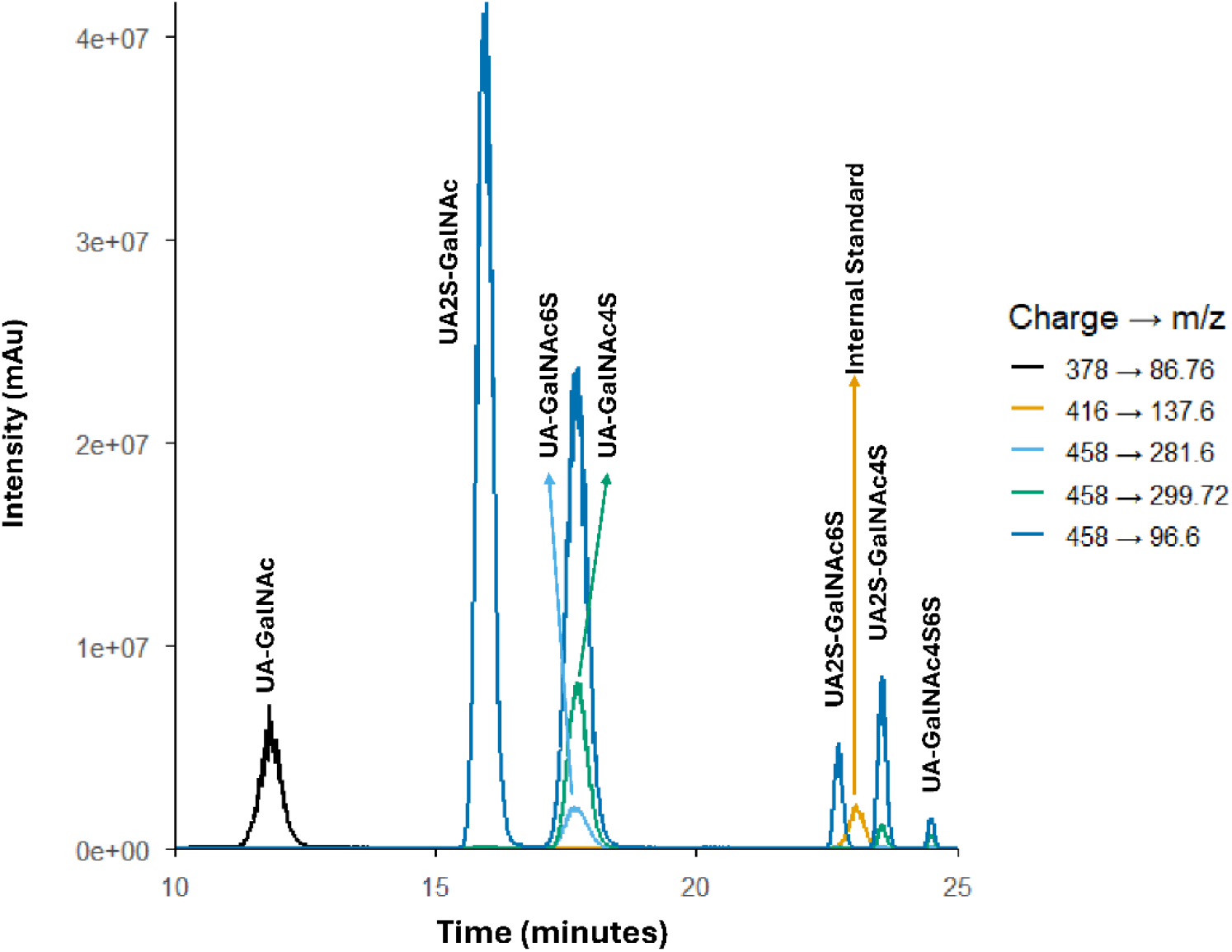
Extracted ion chromatographs of CS standard disaccharides spiked in pooled human milk obtained by LC-MS/MS. Disaccharides were separated by HILIC using a gradient of aqueous ammonium formate (50mM, pH 4.4) and acetonitrile. The MS was operated in negative electrospray ionization mode. Each trace represents a multiple-reaction monitoring transition specific to the disaccharide standards.

Optimal MRM transitions were determined by on-line direct infusion of standards at varying concentrations ranging from 1-10 μg/mL. Precursor ions were identified from full MS scans, and product ion spectra were acquired under collision-induced dissociation (CID) (Supplementary Figures 1-4). For each analyte, multiple transitions were evaluated with final transitions chosen based on signal intensity, fragmentation reproducibility, and absence of matrix interference (Figure 4).

**Figure 4:**
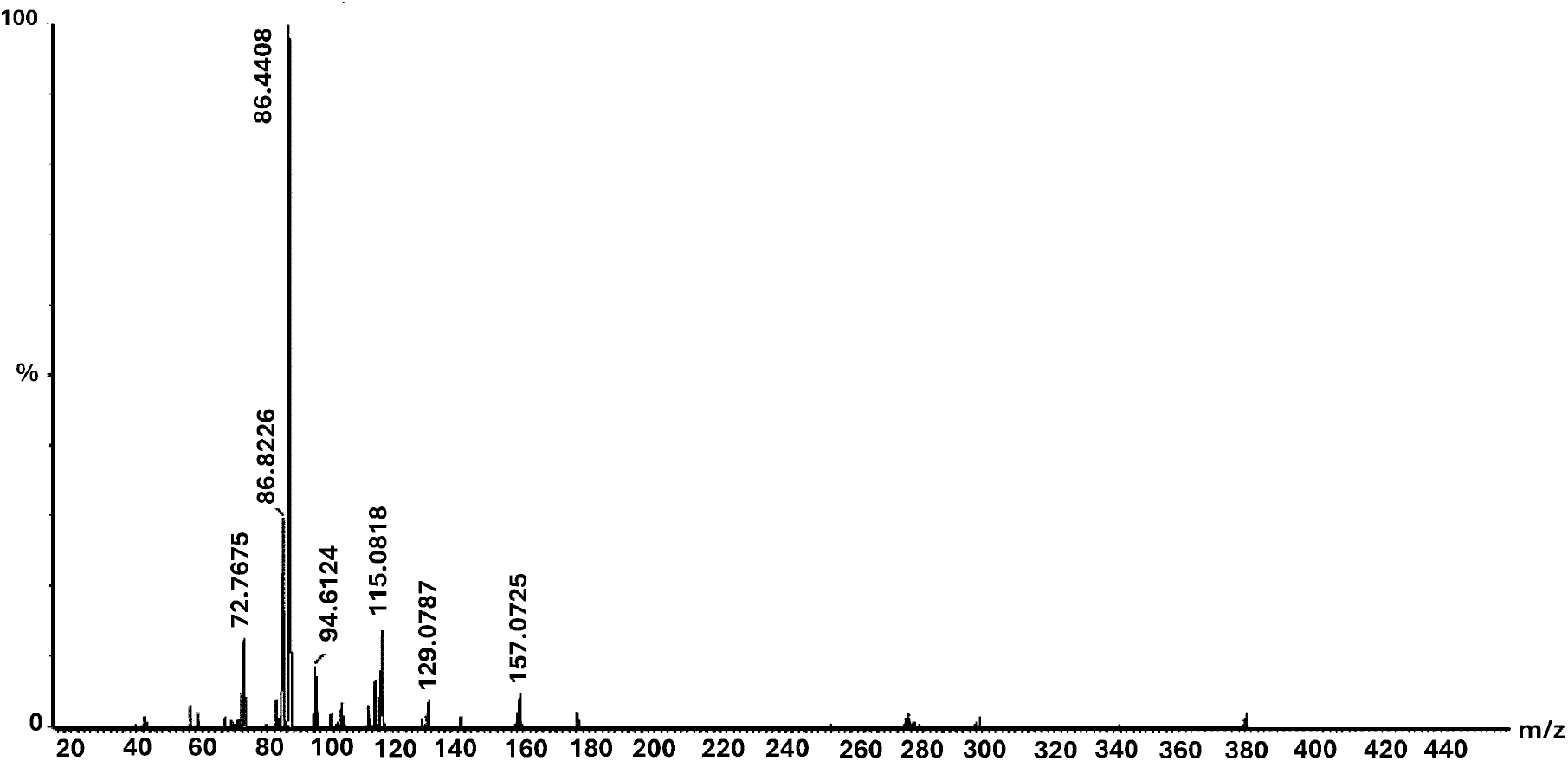
Fragmentation patterns of UA-GalNAc. Fragmentation patterns between 20 – 440 m/z of UA-GalNAc standard after applying collision energy 30 to precursor ion 378.0. Most abundant ion determined as 86.4.

For the isomers UA-GalNAc4S and UA-GalNAc6S, product ion selection focused on identifying transitions capable of distinguishing the two isomers (Figure 5). UA-GalNAc6S m/z 281.74 was the most abundant daughter ion, with minimal presence in UA-GalNAc4S, therefore was selected for detection. Likewise, daughter ion m/z 299.72, with minimal presence in UA-GalNAc6S, was selected for UA-GalNAc4S detection.

**Figure 5:**
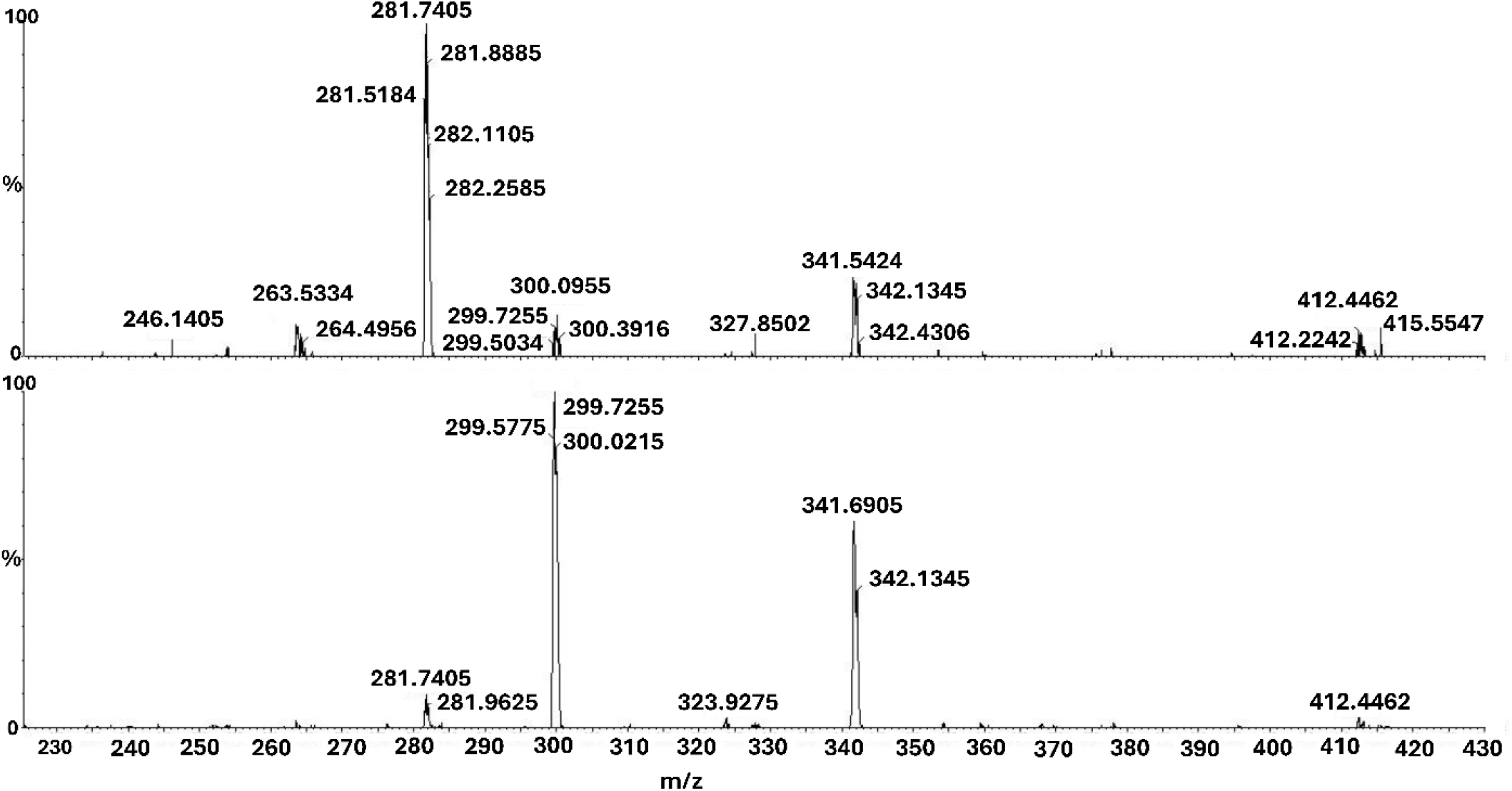
Fragmentation patterns of isomeric pair UA-GalNAc4S and UA-GalNAc6S. Top panel – Fragmentation patterns between 230 – 430 m/z of UA-GalNAc6S standard after applying collision energy 30 to precursor ion 458.0. Bottom panel - Fragmentation patterns between 230 – 430 m/z of UA-GalNAc4S standard after applying collision energy 30 to precursor ion 458.0.

Although the doubly sulphated UA-GalNAc4S6S produced a more intense signal for the 458>96.6 transition corresponding to the loss of a sulphate group, the 458>299.7 transition was ultimately selected (Supplementary Figure 4). This transition exhibited lower background signals in human milk blanks, resulting in improved sensitivity and more stable peak integration across analytical runs.

MS source parameters were optimised to maximise intact ions and MS/MS fragments while minimising in-source sulphate loss. Source parameters were softened (capillary voltage 1.60 kV, cone voltage 23 V) to reduce stress on sulphated disaccharides, and parameters were further tuned with a focus on optimising the signals of internal standard and the disaccharides detected in pooled milk samples (UA‐GalNAc, UA‐GalNAc4S and UA‐GalNAc6S). A relatively high desolvation temperature (600°C) was selected for improved ionization efficiency and signal intensity for the CS disaccharides of interest. Similarly, elevated desolvation gas (1000 L/hr), cone gas (150 L/hr) and nebuliser pressure (7.0 bar) were set to generate a strong, high‐energy drying environment to prevent the strong solvent binding abilities of the highly polar analytes, despite being slightly more aggressive for sulphate retention. This can be seen with the doubly sulphated disaccharides; UA2S-GalNAc4S, UA2S-GalNAc6S and UA-GalNAc4S6S, which are better detected following MRM transitions determined by the loss of one sulphate. MRM transitions for both fully sulphated and loss of a sulphate group were included in the final method to ensure accurate detection. Dwell times were set to ensure approximately 12 data points across each chromatographic peak for peak reproducibility and all final MS settings were verified in matrix‐matched samples.

### Method Validation

#### Linearity, Sensitivity and Model Evaluation

The best calibration model was identified as a weighted (1/x) linear model (not forcing the intercept through 0), supported by the calibration curve plots, response factors and residuals (Figure 6, Supplementary Table 1).

**Table 1.** Mean Signal to Noise (S/N) for CS disaccharides at expected retention time windows in human milk blanks without lyase treatment. S/N was calculated as the mean peak area at each concentration (n=3) divided by the mean background noise measured in non-spiked human milk (matrix blank, n=2).

| ng/mL | Matrix | UA-GalNAc Av. Noise area | UA2S-GalNAc Av. Noise area | UA-GalNAc6S Av. Noise area | UA-GalNAc4S Av. Noise area | UA2S-GalNAc6S Av. Noise area | UA2S-GalNAc4S Av. Noise area | UA-GalNAc4S6S Av. Noise area |
| --- | --- | --- | --- | --- | --- | --- | --- | --- |
| 0 | Non-spiked human milk | 1217661.0 | 94581.5 | 11842.5 | 271837.5 | 155296.0 | 136495.0 | 46966.0 |
| ng/mL | Matrix | UA-GalNAc S/N | UA2S-GalNAc S/N | UA-GalNAc6S S/N | UA-GalNAc4S S/N | UA2S-GalNAc6S S/N | UA2S-GalNAc4S S/N | UA-GalNAc4S6S S/N |
| 100 | Spiked human milk | 2.9 | 122.3 | 190.2 | 20.8 | 3.6 | 7.00 | 1.5 |
| 200 | Spiked human milk | 4.6 | 230.0 | 404.3 | 37.7 | 7.19 | 14.9 | 3.0 |
| 300 | Spiked human milk | 6.2 | 370.9 | 666.8 | 61.1 | 12.55 | 24.7 | 4.3 |
| 400 | Spiked human milk | 7.0 | 468.7 | 843.7 | 78.0 | 15.21 | 31.1 | 5.9 |
| 500 | Spiked human milk | 9.6 | 594.9 | 1109.6 | 103.4 | 18.80 | 40.3 | 7.2 |
| 600 | Spiked human milk | 14.6 | 701.0 | 1306.4 | 120.8 | 22.19 | 47.7 | 8.7 |

**Figure 6:**
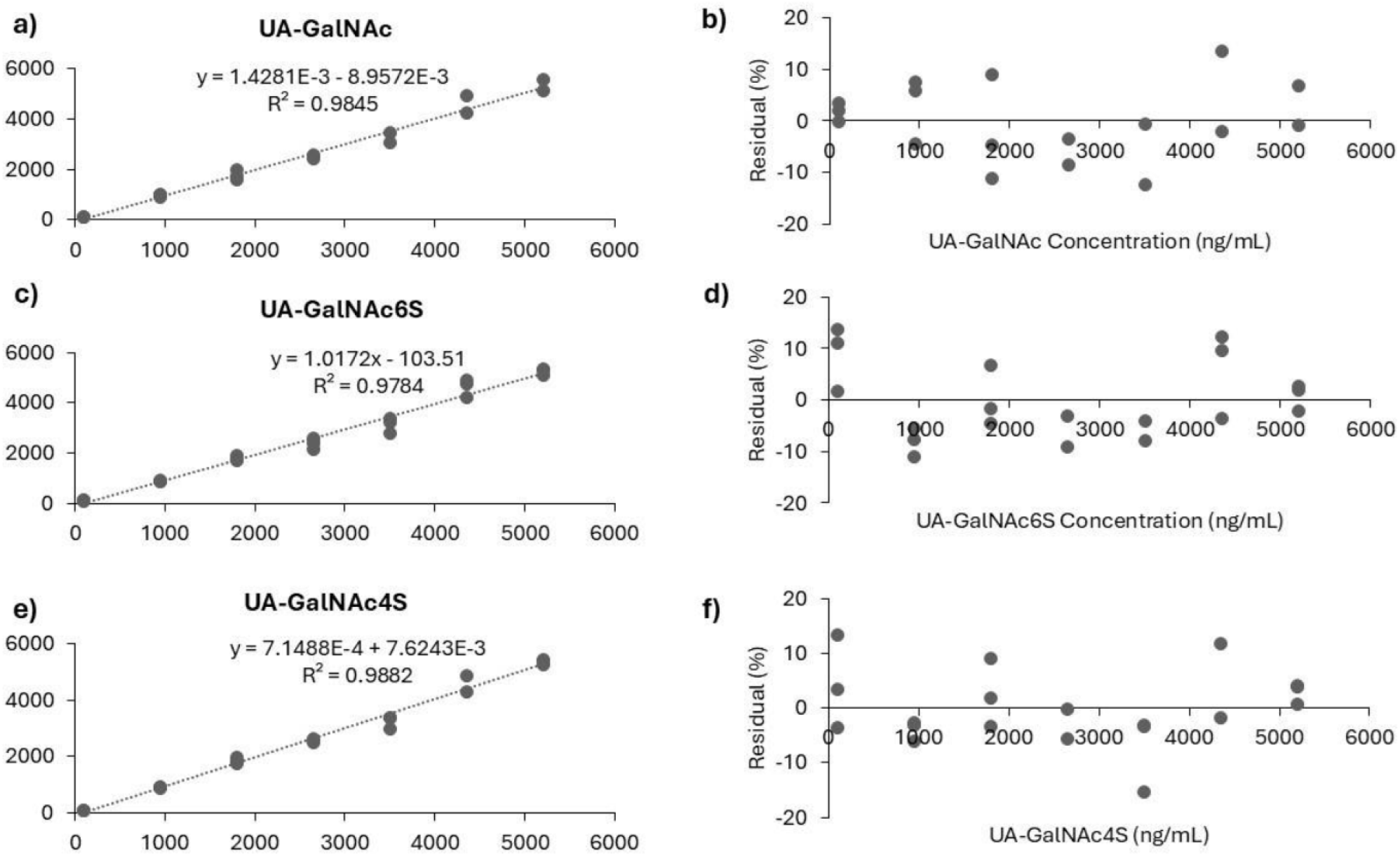
Calibration model of best fit determination. Panel a – Calibration curve for UA-GalNAc. Panel b – Residual plot for linear calibration model of UA-GalNAc. Panel c - Calibration curve for UA-GalNAc6S. Panel d - Residual plot for linear calibration model of UA-GalNAc6S. Panel e - Calibration curve for UA-GalNAc4S. Panel f - Residual plot for linear calibration model of UA-GalNAc4S. Residuals (%) were plotted against concentration (ng/mL) for each analyte across the calibration range with a 1/x weighting.

The weighting of 1/x was found to improve accuracy at low concentrations. Residuals were evenly distributed across the range confirming the appropriateness of the model (Figure 6).

Across the CS disaccharides, clear differences in detectability emerged from the S/N profiles. UA‐GalNAc showed the weakest response, after UA-GalNAc4S6S, with S/N rising from 2.8 at 100 ng/mL to 14.6 at 600 ng/mL, indicating a limit of detection (LoD) just above 100 ng/mL and a limit of quantification (LoQ) near 500–600 ng/mL. In contrast, UA2S‐GalNAc exhibited exceptionally strong signals, with S/N already exceeding 120 at 100 ng/mL and increasing to over 700 at 600 ng/mL; both LoD and LoQ therefore lie well below the lowest calibration point. UA‐GalNAc6S behaved similarly, with S/N values from 190 to over 1300 across the range, again placing both LoD and LoQ below 100 ng/mL. UA‐GalNAc4S also demonstrated high sensitivity, with S/N comfortably above 20 at 100 ng/mL and rising to 121 at 600 ng/mL, supporting an LoQ below 100 ng/mL. UA2S‐GalNAc6S showed moderate detectability, with S/N increasing from 3.6 at 100 ng/mL to 22.2 at 600 ng/mL; this pattern suggests an LoD around 100 ng/mL and an LoQ near 300 ng/mL. UA2S‐GalNAc4S displayed a stronger response, with S/N values from 7.0 to 47.7, placing the LoD below 100 ng/mL and the LoQ in the 200–300 ng/mL range. Finally, UA‐GalNAc4S6S showed the poorest sensitivity, with S/N increasing from 1.5 at 100 ng/mL to 8.7 at 600 ng/mL; this indicates an LoD between 200–300 ng/mL and an LoQ close to 600 ng/mL.

#### Chondroitin Sulphate Baselines in Human Milk

Baseline concentrations were established over seven independent analysis days, with pooled term human milk prepared fresh each day to capture day to day variability in the matrix. The same pooled milk was used for spike recovery experiments, enabling direct comparison between natural background concentrations and baseline-corrected spike responses under identical matrix conditions. Data were evaluated using the AOAC International’s Interlaboratory Study Evaluation Workbook for blind replicates (version 2.1, available on-line at https://www.aoac.org/resources/aoac-international-interlaboaratory-study-workbook-blind-replicates, accessed 11.03.2026). Although the data were not collected as part of a multi-laboratory study, the workbook could still be used to estimate precision as repeatability (RSD_r_) and intermediate precision (RSD_iR_). The tool also includes Grubbs tests to evaluate if any data points behave as statistical outliers. No significant outliers were detected.

The precision data for UA-GalNAc had a repeatability RSD (RSD_r_) of 16.4%, and intermediate precision RSD (RSD_iR_) of 41.6%, indicating substantial day‐to‐day variability. Accordingly, per‐day baseline values were used for spike‐recovery correction. The variability likely reflects matrix effects, including co-eluting compounds that suppress ionization and interfere with accurate detection. Future optimisation may involve incorporating additional sample preparation steps aimed at isolating and removing matrix compounds that impact UA-GalNAc signal intensity and stability. UA-GalNAc6S had an RSD_r_ of 10.6% and RSD_iR_ of 14.8% indicating acceptable intra- and inter-day variability in baselines levels. Global baseline values were calculated for spike-recovery correction. Likewise, UA-GalNAc4S had an RSD_r_ of 9.6% and RSD_iR_ of 17.3%, therefore global baseline values were used for spike-recovery calculations (Table 2).

**Table 2.** Daily baseline concentrations for all CS disaccharides measured in pooled human milk across seven days. Outliers testing was performed using, Grubbs test. *Due to sample preparation issues, day 1 includes a single human milk blank and day 2 includes duplicate human milk blanks. Pooled human milk blanks were ran in triplicate across days 3-8.

| Day | UA-GalNAc<br>Baseline Mean<br>(ng/mL) | UA-GalNAc6S<br>Baseline Mean<br>(ng/mL) | UA-GalNAc4S<br>Baseline Mean<br>(ng/mL) | Total Baseline<br>CS (ng/mL) |
| --- | --- | --- | --- | --- |
| 1* | 395.8 | 101.2 | 378.6 | 946.4 |
| 2 | 479.9 | 111.6 | 439.0 | 688.7 |
| 3 | 243.7 | 119.1 | 325.9 | 569.7 |
| 4 | 188.6 | 92.5 | 288.5 | 572.9 |
| 5 | 157.4 | 96.2 | 320.0 | 760.0 |
| 6 | 283.1 | 116.3 | 360.6 | 547.4 |
| 7 | 181.8 | 89.6 | 275.9 | 688.7 |
| n | 19 | 19 | 19 | 16 |
| Mean | 275.8 | 103.8 | 341.1 | 647.6 |
| $RSD_r$ (%) | 16.4 | 10.6 | 9.6 | 10.9 |
| $RSD_{IR}$ (%) | 41.6 | 14.8 | 17.3 | 19.9 |
| Outliers | None | None | None | None |
| Baseline Strategy | Per-day baseline | Global baseline | Global baseline | - |

Several CS disaccharides (UA2S-GalNAc, UA2S-GalNAc6S, UA2S-GalNAc4S, and UA-GalNAc4S6S) were detectable but exhibited baseline values that fluctuated around zero across the seven days of analysis (Table 3). For each analyte, the day-to-day variability was larger than the mean measured concentration for each analyte, indicating the natural background concentrations as below the methods lower limit of quantification. These disaccharides are detectable below 100 ng/mL, however, not quantifiable (Table 3).

**Table 3.**
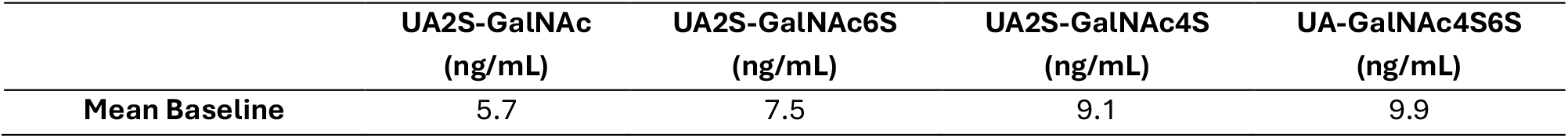
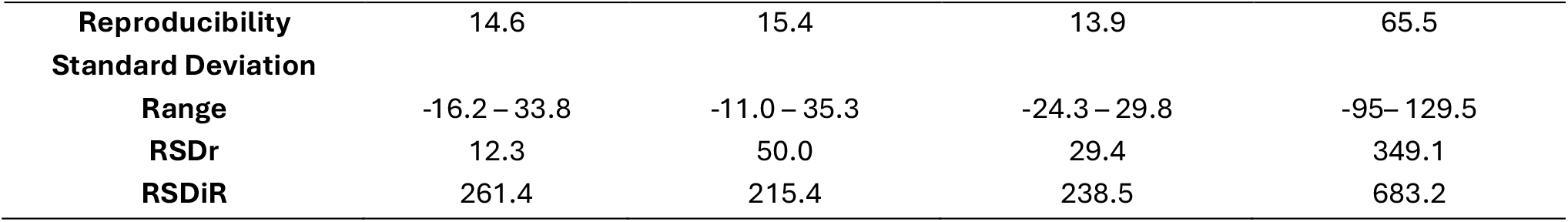
Baseline concentration of low-abundance CS disaccharides in pooled human milk over seven days. Total number of replicates = 8.

#### Accuracy and Precision

Low (1683.8 ng/mL) and high (3302 ng/mL) level spike recoveries were evaluated as a part of the overall method validation. In total, 19 low level and 20 high level spike samples were analysed. Total CS mean recoveries were 41.3% (RSD 22.7%) for the low spikes and 43.7% (RSD 24.6%) for the high spikes (Table 4). Precision was reasonable for the low and non-spiked samples (< 15% RSD_r_, and < 20% RSD_iR_), unusually it was slightly higher for the high spike samples, suggesting there may have been some issues with the spiking. Nevertheless, the data indicate that the method should be well adapted to detect relative differences in CS concentration between samples.

**Table 4.**
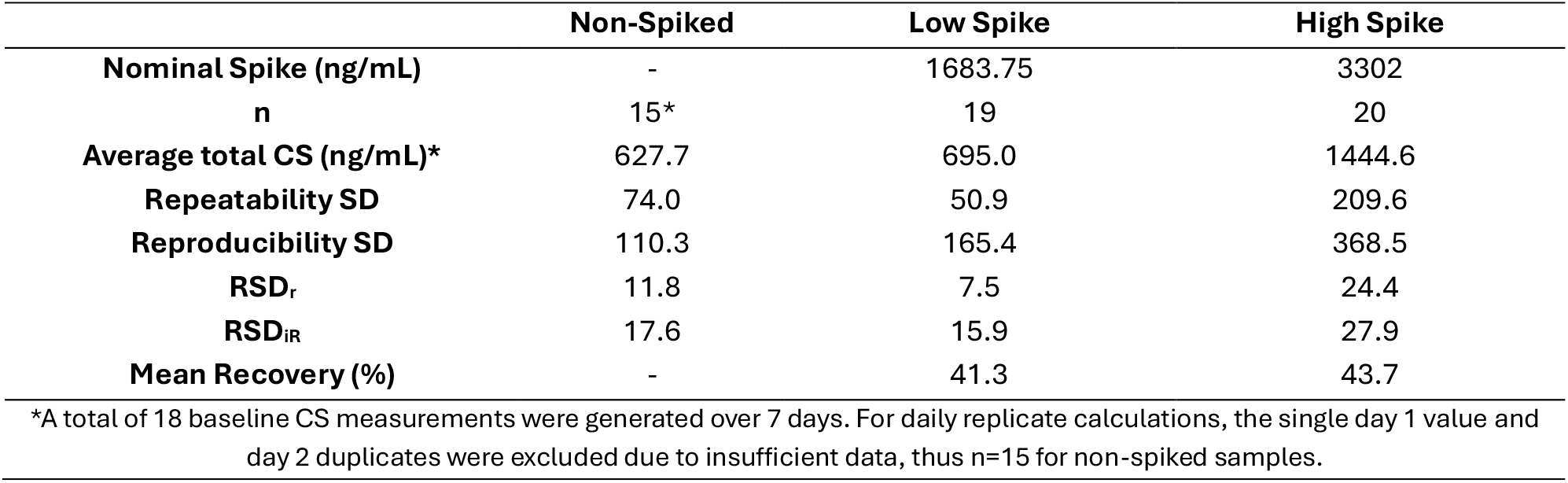
Repeatability, reproducibility, and recovery metrics for baseline, low-spike and high-spike chondroitin sulphate in human milk. Spike values were corrected by baseline signal subtraction.

|  | **Non-Spiked** | **Low Spike** | **High Spike** |
| --- | --- | --- | --- |
| **Nominal Spike (ng/mL)** | - | 1683.75 | 3302 |
| **n** | 15* | 19 | 20 |
| **Average total CS (ng/mL)\*** | 627.7 | 695.0 | 1444.6 |
| **Repeatability SD** | 74.0 | 50.9 | 209.6 |
| **Reproducibility SD** | 110.3 | 165.4 | 368.5 |
| **RSD<sub>r</sub>** | 11.8 | 7.5 | 24.4 |
| **RSD<sub>iR</sub>** | 17.6 | 15.9 | 27.9 |
| **Mean Recovery (%)** | - | 41.3 | 43.7 |

\*A total of 18 baseline CS measurements were generated over 7 days. For daily replicate calculations, the single day 1 value and day 2 duplicates were excluded due to insufficient data, thus n=15 for non-spiked samples.

#### Outlooks

Understanding human milk composition is a crucial aspect to understanding infant health. This method offers several important advantages, particularly the ability to work with much smaller sample volumes (25 μL) providing access to samples previously considered unusable for GAG determination in human milk, enabling meaningful analysis with limited material. This can be especially valuable in longitudinal studies across lactation of ‘endangered’ cohorts of infants where only small aliquots may be available, and high-volume ‘gold-standard’ collection approaches are not feasible. A drawback to the small sample volume is increased variability due to imprecisions of manipulating small volumes or weights of sample and reagents. However, this does not necessarily indicate that the lower recoveries were caused by reduced sample volume; rather, it is likely that loss of recovery is associated with multiple sample preparation steps.

Although absolute recoveries did not reach the desired performance thresholds, they were consistent at the two spike levels investigated during validation, demonstrating suitability of the method to compare CS levels in different milk samples (Table 4). Moreover, this approach minimises opportunities for variability to be introduced by prolonged storage and extended sample handling over multiple days. This method provides clear opportunities for optimisation. For example, the incorporation of a β‐galactosidase and proteinase treatment may reduce background signal throughout chromatography and mitigate interference from compounds that could interfere with recoveries.

Recovery experiments performed in neat aqueous solution (CS-A in water) using a single fixed calibration curve across 8 days yielded mean recoveries of 71% (low spike) and 60% (high spike). Alternatively, CS-A spiked in human milk and quantified using a fresh calibration curve prepared daily, achieved recoveries of 41.3% for the low spike, and 43.7% for the high spike. This reduction in recovery in human milk demonstrates a pronounced matrix effect, consistent with ion suppression and/or incomplete extraction in this complex biological matrix.

## CONCLUSION

A novel method has been developed for the determination of CS in small volumes (25 μL) of human milk which also enables the estimation of the relative ratios of 4- and 6-sulfation. This method provides the foundation for absolute CS quantification with further optimisation. In the interim, this method allows for the comparative analysis of CS in low volumes of human milk (which was not possible with existing protocols). The streamlined sample preparation protocol facilitates same day preparation and analysis of samples. This protocol enables CS profiling in human milk from minimal sample volumes, expanding the feasibility of studying human milk composition across diverse cohorts.

## MATERIALS AND METHODS

### Standards and Reagents

Chondroitinase ABC from Proteus vulgaris, Chondroitin sulphate A, Acetonitrile, Bovine serum albumin, Ammonium Formate, TRIS Base Ultra-Pure, and Calcium chloride were obtained from Merck, Gillingham, UK. Unsaturated CS disaccharides were obtained from Iduron, Alderley Edge, UK. Water (LC-MS, UPLC-UV Grade), Hydrochloric Acid was obtained from Fisher Scientific, Loughborough, UK. ‘Breast Milk (Pooled from 3 Human Donors delivering term infants) was obtained from Lee BioSolutions, Inc, Maryland Heights, MO USA 63043.

#### Reagent Preparation

Chondroitinase ABC stock solution was prepared in aliquots by dissolving chondroitinase ABC (10U) in 0.01% of BSA to a final concentration of 2 U/mL. Aliquots (100 μL) were stored at −20°C in microtubes. On the day of analysis, chondroitinase ABC stock solution was thawed and further diluted with 0.01% BSA to a final concentration of 0.2 U/mL.

The LC aqueous eluent (50mM Ammonium Formate, pH 4.4) was prepared fresh each week by dissolving the reagent into MS grade water and sonicating for 5 minutes, prior to pH adjustment using HCl.

Precipitation buffer (15/85_(v/v)_ Ammonium Formate (50mM)/Acetonitrile_(v/v)_) was prepared by combining 15 mL of the LC aqueous buffer with 8 mL MS grade acetonitrile and inverting three times prior to use.

CS-A spike stock solutions were prepared in one batch for use across the entire validation study by preparing volumes of CS-A spike material in water at 1682.8 ng/mL and 3302 ng/mL. Spike material in water was then combined in equal parts with the pooled human milk _(v/v)_ and aliquoted for daily use prior to storage at −20°C. This approach ensured all calibration and QC materials were prepared from a single batch of matrix-matched spike material, minimising day-to-day variability associated with repeated spiking, and reduced freeze-thaw cycles of the stock solution.

#### Standards preparation

Quantification of CS was performed by preparing the ‘CS Curve Mix’ at a concentration of 10 μg/mL using stocks of ΔUA-GalNAc, ΔUA2S-GalNAc, ΔUA-GalNAc6S, ΔUA-GalNAc4S, ΔUA2S-GalNAc6S, ΔUA2S-GalNAc4S, and ΔUA-GalNAc4S6S. Using the CS Curve Mix, seven calibration curve points were prepared at 100 ng/mL, 950 ng/mL, 1800 ng/mL, 2650 ng/mL, 3500 ng/mL, 4350 ng/mL, and 5200 ng/mL.

### Equipment

ACQUITY™ Premier Glycan BEH Amide, 130Å, 1.7μm 2.1 × 150mm Column, Xevo TQ-XS Triple Quadrupole Mass Spectrometer, Acquity UPLC I-Class PLUS System were obtained from Waters Corporation (Wilmslow, Cheshire, UK). SevenCompact pH meter was obtained from Mettler-Toledo Ltd. (Leicester, UK). 20-200 μL micropipettes and Benchtop Mixer HC (1.5 mL) Thermoshaker were obtained from Starlab (UK). Benchtop sonic bath and fixed insert (300 μL) amber LC vials obtained from Thermo Scientific (USA). Vortex mixer from IKA (Germany), benchtop centrifuge from Sciquip (UK), microcentrifuge tubes (1.5 mL) obtained from Eppendorf (UK). Benchtop scales (SECURA225D-1S) obtained from Sartorius AG.

### Protocol to Determine CS in Human Milk

CS standard curve was prepared as detailed in standards preparation.

Prior to analysis, frozen samples were thawed at room temperature and mixed well by vortexing. The calibration curve in matrix, control human milk samples, CS-A spiked human milk samples and CS-A spiked water samples were prepared following the procedure in Table 5.

**Table 5.** Sample preparation protocol details. All spike samples were prepared at low and high spiked in triplicate.

| Calibration Curve | Control Breastmilk | CS-A Spiked Breastmilk (Med/High) | CS-A Spiked Water |
| --- | --- | --- | --- |
| 25 µL Human milk | 25 µL Human milk | 50 µL Spiked Human milk | 25 µL CS-A Spike Material in Water |
| 25 µL Curve mix | 25 µL MS MiliQ Water |  | 25 µL MS MiliQ Water |
| 25 µL Enzyme Working Solution<sup>1</sup> | 25 µL Enzyme Working Solution<sup>1</sup> | 25 µL Enzyme Working Solution<sup>1</sup> | 25 µL Enzyme Working Solution<sup>1</sup> |
| 25 µL BSA (0.01%) (No Enzyme)<sup>2</sup> | 25 µL chondroitinase ABC in BSA | 25 µL chondroitinase ABC in BSA | 25 µL chondroitinase ABC in BSA |

<sup>1</sup> Enzyme Working Solution (EWS) = 50mM TRIS, 5mM CaCl<sub>2</sub>, pH 8.1 containing internal standard at 2650 ng/mL. <sup>2</sup> Calibration curve samples contain no enzyme.

All samples were then incubated for 5 hours at 37°C, 16162 × g using the Thermomixer, then centrifuged at 16162 × g for 30 minutes prior to refrigeration for 15 minutes. 80 μL of sample was then transferred to a fresh Eppendorf, avoiding the top fat layer and lower pellet and 452 μL of precipitation buffer was added. Samples were then centrifuged for 15 minutes at 16162 × g, prior to another round of refrigeration. For the analysis of samples, 250 μL of the supernatant was transferred to HPLC vials, avoiding any precipitate at the bottom of the Eppendorf tubes.

### Liquid Chromatography Mass Spectrometry Conditions

#### Chromatographic Conditions

Analysis was performed using an Acquity UPLC I-Class PLUS System coupled to a Xevo TQ-XS Triple Quadrupole Mass Spectrometer operated *via* the MassLynx software. Chromatographic separation was achieved on an ACQUITY™ Premier Glycan column (130Å, 1.7μm 2.1 × 150mm) at 80°C and an eluent pre-heater also set at 80°C. Analytes were eluted using a linear gradient of 50mM ammonium formate, pH 4.4 in acetonitrile (Table 6). Injection volume was set at 10 μL. The reduction in flow rate from between 32.1 and 38.0 minutes prevented system overpressure during the column wash and re-equilibration steps.

**Table 6.** HILIC Gradient for separation of CS/DS disaccharides.

| Time<br>(min) | Flow (mL/min) | %A<br>(Acetonitrile) | %B<br>(Ammonium<br>formate, 50<br>mM, pH 4.4) |
| --- | --- | --- | --- |
| 2.0 | 0.600 | 88.0 | 12.0 |
| 9.0 | 0.600 | 85.0 | 15.0 |
| 15.0 | 0.600 | 82.0 | 15.0 |
| 19.0 | 0.600 | 82.0 | 18.0 |
| 24.0 | 0.600 | 72.6 | 27.4 |
| 32.0 | 0.600 | 54.0 | 46.0 |
| 32.10 | 0.300 | 30.0 | 70.0 |
| 37.0 | 0.300 | 30.0 | 70.0 |
| 38.0 | 0.300 | 88.0 | 12.0 |
| 40.0 | 0.600 | 88.0 | 12.0 |
| 48.0 | 0.600 | 88.0 | 12.0 |

#### MS Parameters

The MS analyses were performed using ESI in negative mode with capillary voltage at 1.60 kV and cone voltage at 23 V. Desolvation temperature was set to 600°C, with Desolvation gas 1000 L/Hr, Cone gas 150 L/Hr and Nebuliser gas 7.0 Bar. Collision gas flow was set to 0.15 mL/min with a resting collision energy of 4 V to enhance pre-cursor detection. The MRM transitions monitored are presented in Table 7.

**Table 7.**
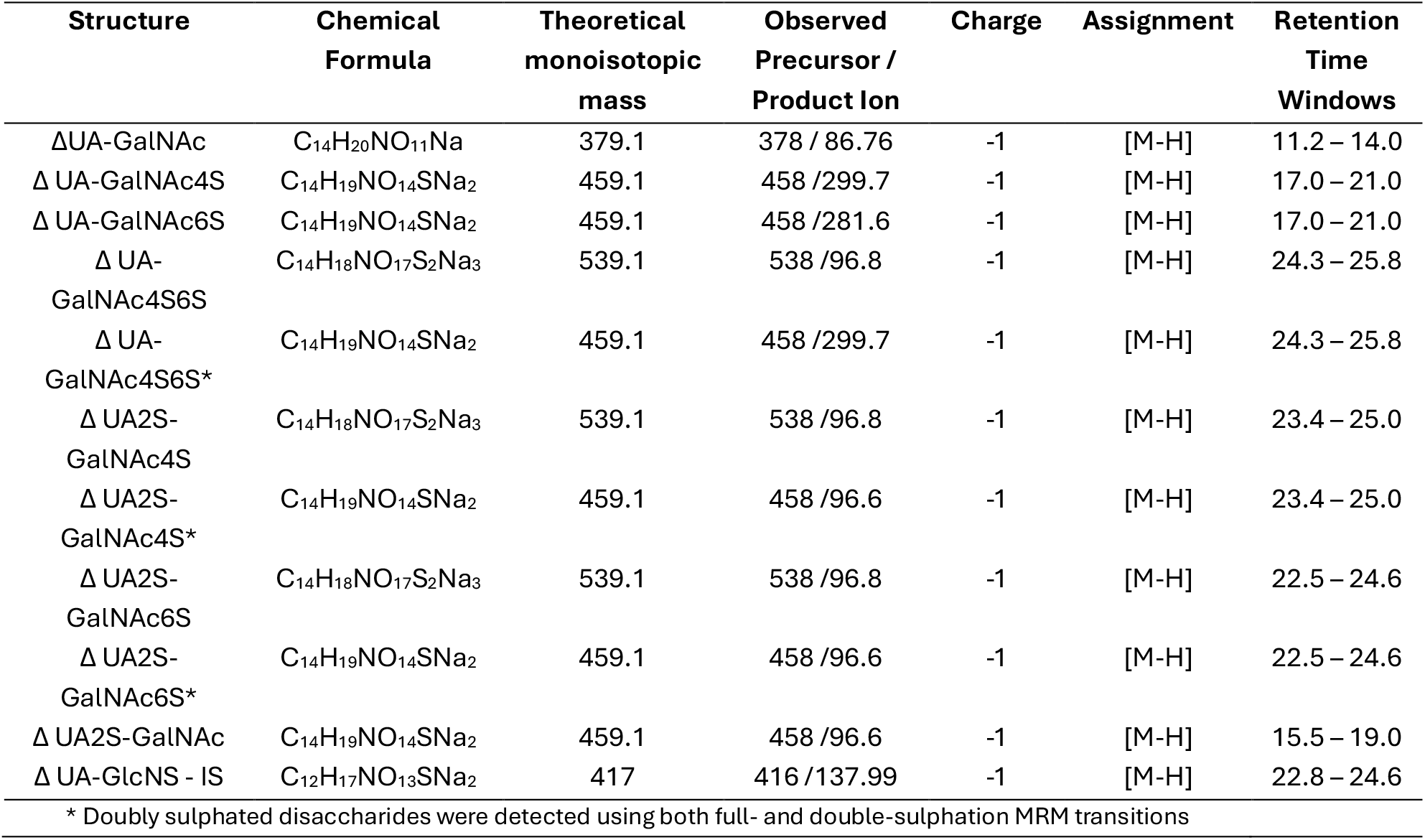
MRM transitions for CS disaccharides. Retention time windows are large to account for shape of HILIC produced peaks. Slight retention time window shift is observed across runs.

### Method Validation Design

#### Calibration Model

Assessment of calibration model was performed using a seven-point CS disaccharide calibration range (100–5200 ng/mL). On a single day, 21 calibration solutions representing three independent calibration curves were prepared, and each solution was analysed once. This approach allowed us to evaluate the reproducibility of curve preparation and to determine the most appropriate regression model and weighting strategy. Model fit was assessed using the coefficient of determination (R^2^) and plots of residuals.

#### Limit of Detection and Limit of Quantification

LoD and LoQ were determined using pooled human milk without lyase treatment as the blank matrix. Two non-spiked milk samples were processed identically to the calibration curve to determine baseline noise at the expected retention time windows for each disaccharide. The baseline noise within the expected retention time windows of each analyte was assessed on the samples. In parallel, a six-point CS calibration curve (100-600 ng/mL) prepared in the pooled milk was analysed to determine low-level signal response. S/N ratios were calculated as the mean peak area at each concentration divided by the mean background noise measured in the blank milk (Table 1). e. LoD and LoQ were defined as the concentrations producing signals threefold and tenfold above baseline noise, respectively.

#### Assessment of Method Trueness and Precision

Accuray and precision were assessed by spike-recovery experiments. Low (1683.8 ng/mL) and high (3302 ng/mL) level spike recoveries, using pooled human milk, were evaluated as a part of the overall method validation, as well as human milk analysed without spike material. CS-A spikes (in water) were prepared fresh each day in triplicate. CS-A was chosen as the spike material because it contains the panel of expected CS disaccharides in human milk.

For each day of analysis, freshly prepared spiked pooled human milk, at both the low and high level were prepared and analysed in triplicate, generating multiple spike replicates across the seven-day study.

Precision was evaluated using repeatability and intermediate reproducibility statistics produced from the data gathered across the spike-recovery experiments. Intra-day precision was assessed as the RSD_r_, based on the variability of triplicate measurements within a single day. Inter-day precision was assessed as the RSD_iR_, calculated from the variability of the measurements across seven days.

## ABBREVIATIONS

2-AB: 2-Aminobenzamide
2-AMAC: 2-Aminoacridone
BSA: Bovine Serum Albumin
CID: Collision Induced Dissociation
CS: Chondroitin Sulphate
DS: Dermatan Sulphate
EWS: Enzyme Working Solution
FLD: Fluorescence Detection
GAGs: Glycosaminoglycans
HCl: Hydrochloric Acid
HILIC: Hydrophilic Interaction Liquid Chromatography
IS: Internal Standard
LC: Liquid Chromatography
LoD: Limit of Detection
LoQ: Limit of Quantification
MRM: Multi-Reaction Monitoring
MS: Mass Spectrometry
RP-HPLC-ESI: Reverse Phase High Performance Liquid Chromatography Electrospray Ionization
RSD_iR_: Relative standard deviation in intermediate reproducibility conditions
RSR_r_: Relative standard deviation under repeatability conditions
RSD: relative standard deviation
S/N: Signal to Noise
UPLC: Ultra Performance Liquid Chromatography.

## ACKNOWLEDGEMENTS

MEG is funded by a BBSRC collaborative training partnership PhD studentship. CJS is supported by the Sir Henry Dale Fellowship jointly funded by the Wellcome Trust and the Royal Society (Grant Number 221745/Z/20/Z and the 2021 Lister Institute Prize Fellow Award. The authors would like to thank the Faculty of Science and Engineering Mass Spectrometry and Separation Facility (RRID: SCR_024761).

## DATA AVAILABILITY STATEMENT

Data supporting the findings of this study are available from the corresponding author upon reasonable request.

