## Supplementary Material for "Label-Free Determination of Chondroitin Sulphate from Microgram Quantities of Human Milk"

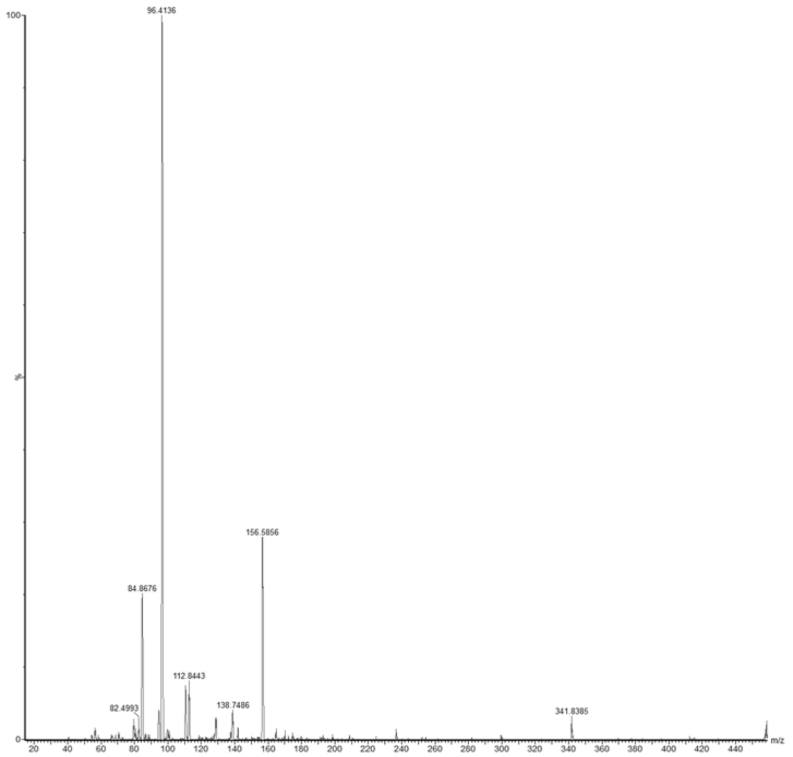


**Supplementary Figure 1:** Fragmentation patterns between 20 – 440 m/z of UA2S-GalNAc standard after applying collision energy 30 to precursor ion 458.0. Most abundant ion range 96.3 – 96.7m/z, followed by second transition ion ~156.0.

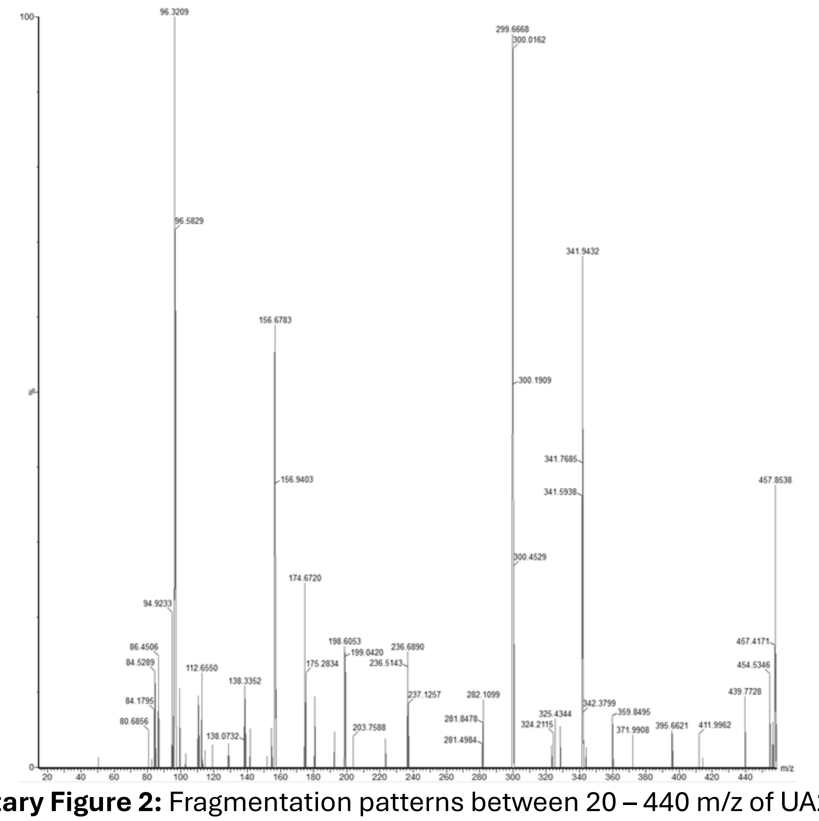


**Supplementary Figure 2:** Fragmentation patterns between 20 – 440 m/z of UA2S-GalNAc4S standard after applying collision energy 30 to precursor ion 538.0. Most abundant ion range 96.3 – 96.7m/z. Product ion 299.7 is equally traceable.

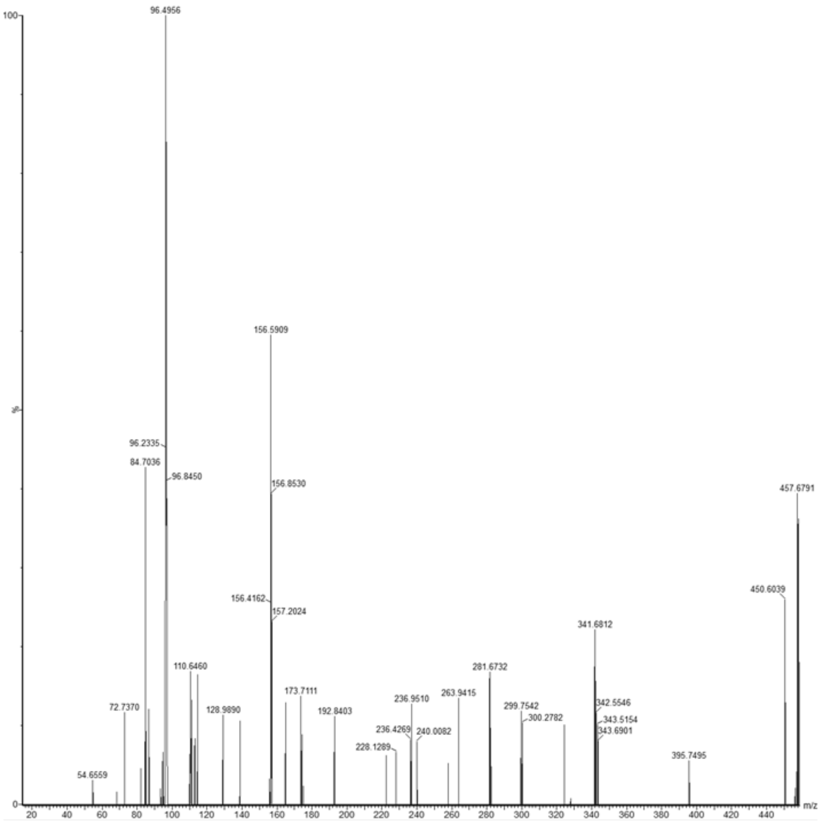


**Supplementary Figure 3:** Fragmentation patterns between 20 – 440 m/z of UA2S-GalNAc6S standard after applying collision energy 30 to precursor ion 538.0. Most abundant ion range 96.3 – 96.7m/z. Secondary transition ions ~156.6 and ~281.6 are abundant transitions.

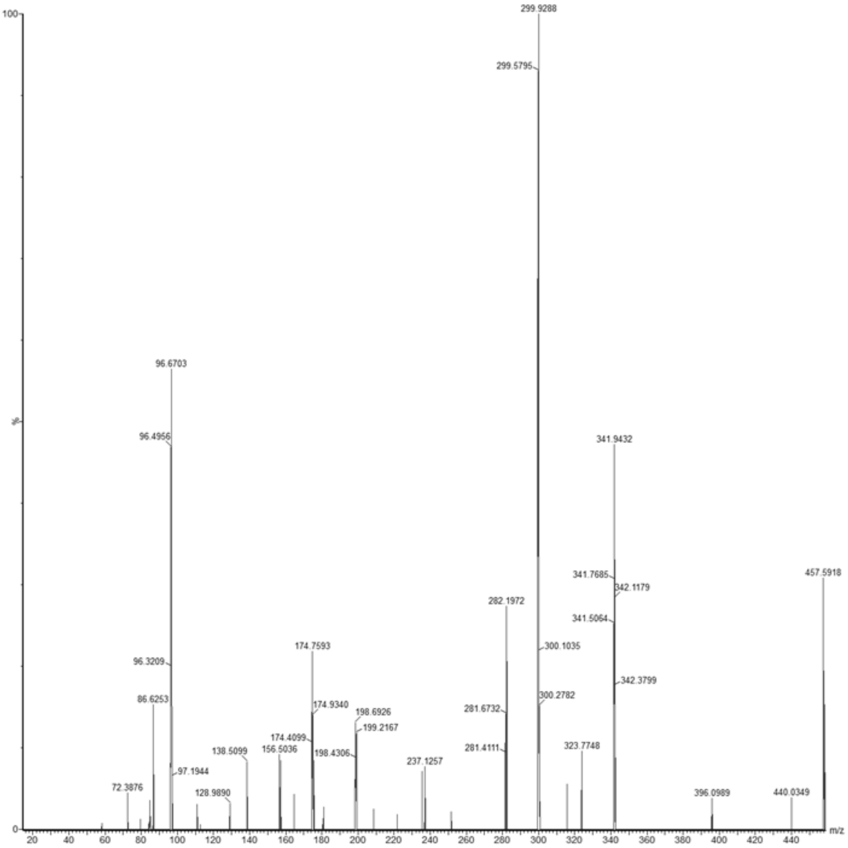

**Supplementary Figure 4:** Fragmentation patterns between 20 – 440 m/z of UA-GalNAc4S6S standard after applying collision energy 30 to precursor ion 538.0. Most abundant ion range 299.5 – 299.9. Secondary transition ions ~96.6 and ~174.7 can also be followed if required.

**Supplementary Table 1:** Daily calibration linearity ($R^2$) for each CS disaccharide across seven independent validation days. All curve points were within ±15% of the expected concentration. All curves had to contain >75% of the original curve points to be valid.

| | **UA-GalNAc $R^2$** | **UA2S-GalNAc $R^2$** | **UA-GalNAc6S $R^2$** | **UA-GalNAc4S $R^2$** | **UA2S-GalNAc6S $R^2$** | **UA2S-GalNAc4S $R^2$** | **UA-GalNAc4S6S $R^2$** |
|---|---|---|---|---|---|---|---|
| Day 1 | 0.999 | 0.9985 | 0.9929 | 0.9912 | 0.9966 | 0.9992 | 0.996 |
| Day 2 | 0.99623 | 0.995 | 0.9986 | 0.9968 | 0.9913 | 0.99 | 0.9982 |
| Day 3 | 0.9953 | 0.991 | 0.9969 | 0.9907 | 0.9986 | 0.9919 | 0.9901 |
| Day 4 | 0.9868 | 0.9949 | 0.9979 | 0.9979 | 0.9985 | 0.991 | 0.9982 |
| Day 5 | 0.9955 | 0.9982 | 0.9944 | 0.9962 | 0.9994 | 0.9986 | 0.9961 |
| Day 6 | 0.9913 | 0.9914 | 0.9992 | 0.9992 | 0.9878 | 0.9853 | 0.9951 |
| Day 7 | 0.9961 | 0.9944 | 0.9894 | 0.9859 | 0.9976 | 0.9968 | 0.9914 |
| Mean $R^2$ | 0.9943 | 0.9945 | 0.9956 | 0.9940 | 0.9957 | 0.9933 | 0.99501 |
